# Synaptic combinatorial molecular mechanisms generate repertoires of innate and learned behavior

**DOI:** 10.1101/500389

**Authors:** Noboru H. Komiyama, Louie N. van de Lagemaat, Lianne E. Stanford, Charles M. Pettit, Douglas J. Strathdee, Karen E. Strathdee, David G. Fricker, Eleanor J. Tuck, Kathryn A. Elsegood, Tomás J. Ryan, Jess Nithianantharajah, Nathan G. Skene, Mike D. R. Croning, Seth G. N. Grant

## Abstract

Although molecular mechanisms underpinning specific behaviors have been described, whether there are mechanisms that orchestrate a behavioral repertoire is unknown. To test if the postsynaptic proteome of excitatory synapses could impart such a mechanism we conducted the largest genetic study of mammalian synapses yet undertaken. A repertoire of sixteen innate and learned behaviors was assessed from 290,850 measures in 55 lines of mutant mice carrying targeted mutations in the principal classes of postsynaptic proteins. Each innate and learned behavior used a different combination of proteins. These combinations were comprised of proteins that amplified or attenuated the magnitude of each behavioral response. All behaviors required proteins found in PSD95 supercomplexes. We show the vertebrate increase in proteome complexity drove an expansion in behavioral repertoires and generated susceptibility to a wide range of diseases. Our results reveal a molecular mechanism that generates a versatile and complex behavioral repertoire that is central to human behavioral disorders.

## Introduction

Animals employ a diverse repertoire of behavioral responses to varied and changing environments. 19^th^ century naturalists and psychologists recognized that the behavioral repertoire of animals had an organization or structure in which elementary behaviors, such as reflexes, were basic building blocks for more complex instincts, higher forms of cognition and ultimately consciousness^1–6^. The basic building blocks of the behavioral repertoire were further classified into innate (inherited) or learned responses^7^. Darwin and Tinbergen also recognized the crucial importance of the appropriate magnitude of any behavioral response^3,8^. They considered this a key factor in the adaptation of an animal to its environmental niche. Inappropriately tuned behavioral responses in humans are the hallmarks of the maladaptive behaviors that characterize developmental, neurological and psychiatric disorders.

No molecular mechanism has yet been described that explains the organization and tuning of the behavioral repertoire, its evolution, or role in human behavioral disorders. Current models for the neurobiological substrates of the behavioral repertoire largely rest on the view that different brain regions and circuits underpin individual behaviors. Although molecular and genetic studies have identified genes involved with individual behaviors, whether there are any molecular mechanisms that might regulate multiple elementary behaviors and thereby control a behavioral repertoire is unknown. A precedent for a biological mechanism controlling a repertoire of responses to diverse environmental challenges is found in the combinatorial molecular mechanisms of the vertebrate immune system^9^.

To search for molecular mechanisms of a mammalian behavioral repertoire, we focused on the postsynaptic terminal of excitatory synapses for the following reasons. First, excitatory synapses are components of most brain circuits and behaviors. Second, the postsynaptic proteome of excitatory synapses is highly complex, with many classes of proteins assembled into signaling complexes and networks^10–19^. Third, there is a remarkable number of behavioral disorders that arise from mutations in genes encoding the excitatory postsynaptic proteome. These include over 130 monogenic and polygenic brain disorders, including schizophrenia, depression, autism and intellectual disability^11,20–25^. Mutations in genes encoding postsynaptic excitatory proteins are responsible for 68% of the 94 most common *de novo* mutations causing developmental disorders^26^. Finally, although there have been many mouse genetic studies of postsynaptic proteins, those studies have typically measured the phenotype of an individual gene in specific behaviors rather than examining a repertoire of behavior, nor have they compared the phenotypes of a large panel of genes side-by-side. Thus, in addition to studying the biological basis of the behavioral repertoire, a large-scale genetic dissection of the postsynaptic proteome would provide valuable information on the basic function of excitatory synapses and potentially explain why so many behavioral disorders are caused by mutations disrupting postsynaptic proteins.

We chose to measure a set of innate and learned behaviors using standard test apparatus including the elevated plus maze, open field, novel object exposure task, a rotating rod and a chamber for cue and contextual fear conditioning. The innate behavioral responses measured in these apparatus are frequently described as locomotion, exploration, curiosity, novelty seeking, anxiety and defense responses^27^. We also measured several forms of learning and memory in order to assess whether their genetic mechanisms were distinct from those regulating innate behaviors. In addition to designing a large-scale behavioral experiment, we also measured synaptic physiology^28^ and transcriptomes in the hippocampus. These datasets entailed 290,850 behavioral measures in 2,770 mice, 51,048 measures of electrophysiology in 2,836 hippocampal slices, and genome-wide transcriptome analysis on 390 mice, representing the largest genetic study of mammalian synapses to date. A key part of our strategy was to adopt standardized protocols, which permitted us to quantify and compare massive datasets for many genes and apply robust statistical methods. All datasets are freely available at the Genes to Cognition (G2C) Programme database http://www.genes2cognition.org/publications/g2c/. Here, we report the genetic dissection of the behavioral repertoire and in a companion manuscript we report the genetic dissection of synaptic electrophysiology and integration with behavioral data^28^.

## Results

### The Genes to Cognition phenotyping pipeline

We constructed a highly standardized pipeline of mouse mutant construction, breeding, housing and phenotyping within the Wellcome Trust Sanger Institute animal facility (Fig. 1). The behavior of 55 lines of mice carrying mutations in 51 genes was examined (Fig. 1, 2B, Supplementary Data 1, http://www.genes2cognition.org/publications/g2c/). Several lines were previously characterized and served as a benchmark for our experiments (a comparison of our results with published studies is shown in Supplementary Table 1). Most of the genes were previously uncharacterized and chosen because they encoded seven representative classes of postsynaptic proteins described in proteomic studies of the postsynaptic density (PSD): glutamate receptors, surface adhesion proteins, adaptors and scaffolders, structural proteins, small G-proteins and regulators, signaling enzymes, kinases and phosphatases (Fig. 2B, 3). Fifty lines harbored loss-of-function (LoF) mutations (homozygous null or heterozygous haploinsufficient) and another five lines carried knockin mutations (Fig. 3). The gene set included phylogenetically conserved synapse proteins, paralogs from six protein families and orthologs of human disease genes.

**Figure 1.**
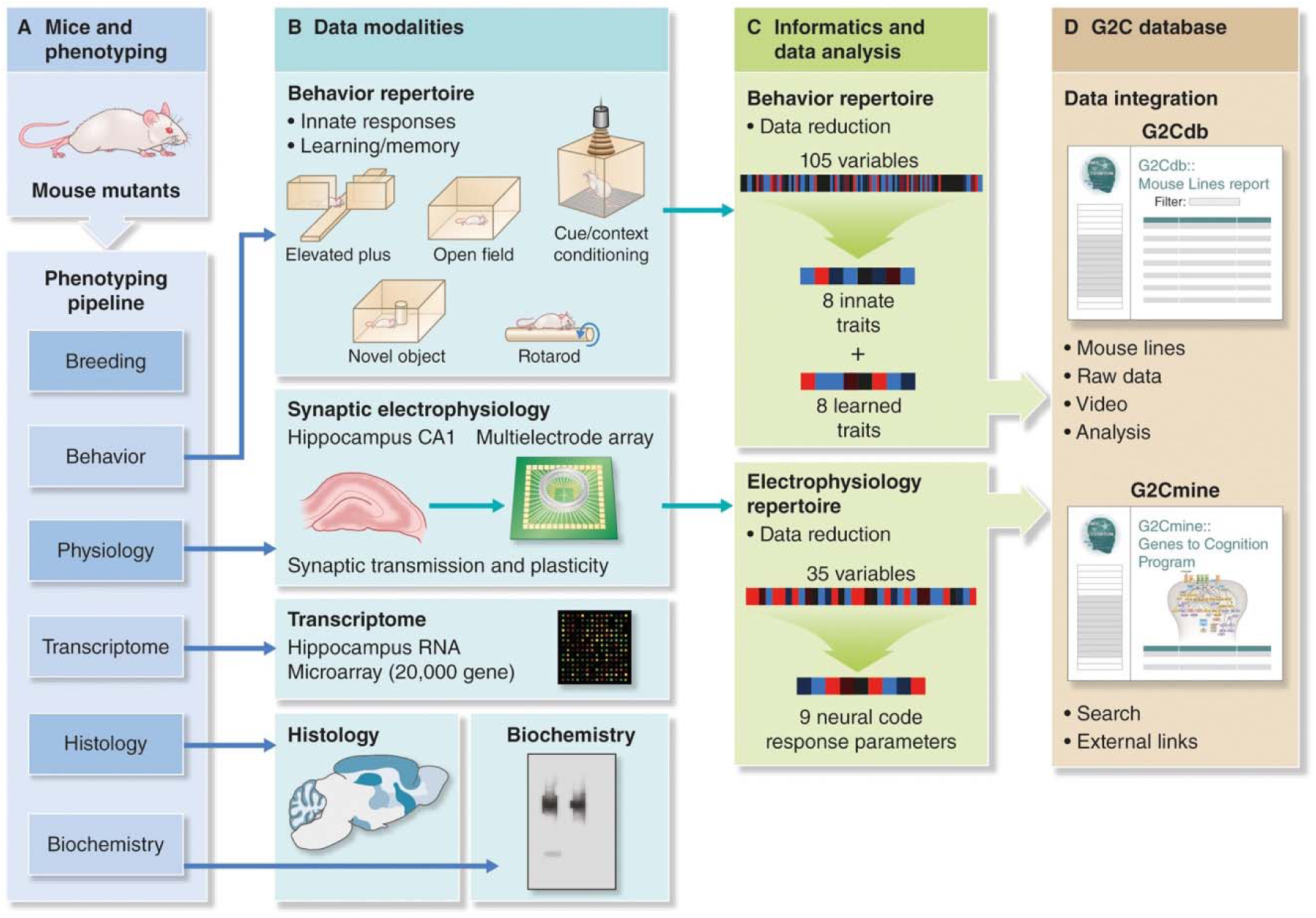
Genes to Cognition Program. Workflow of mutant mouse generation, standardized data acquisition, bioinformatics analysis and data integration.

**Figure 2.**
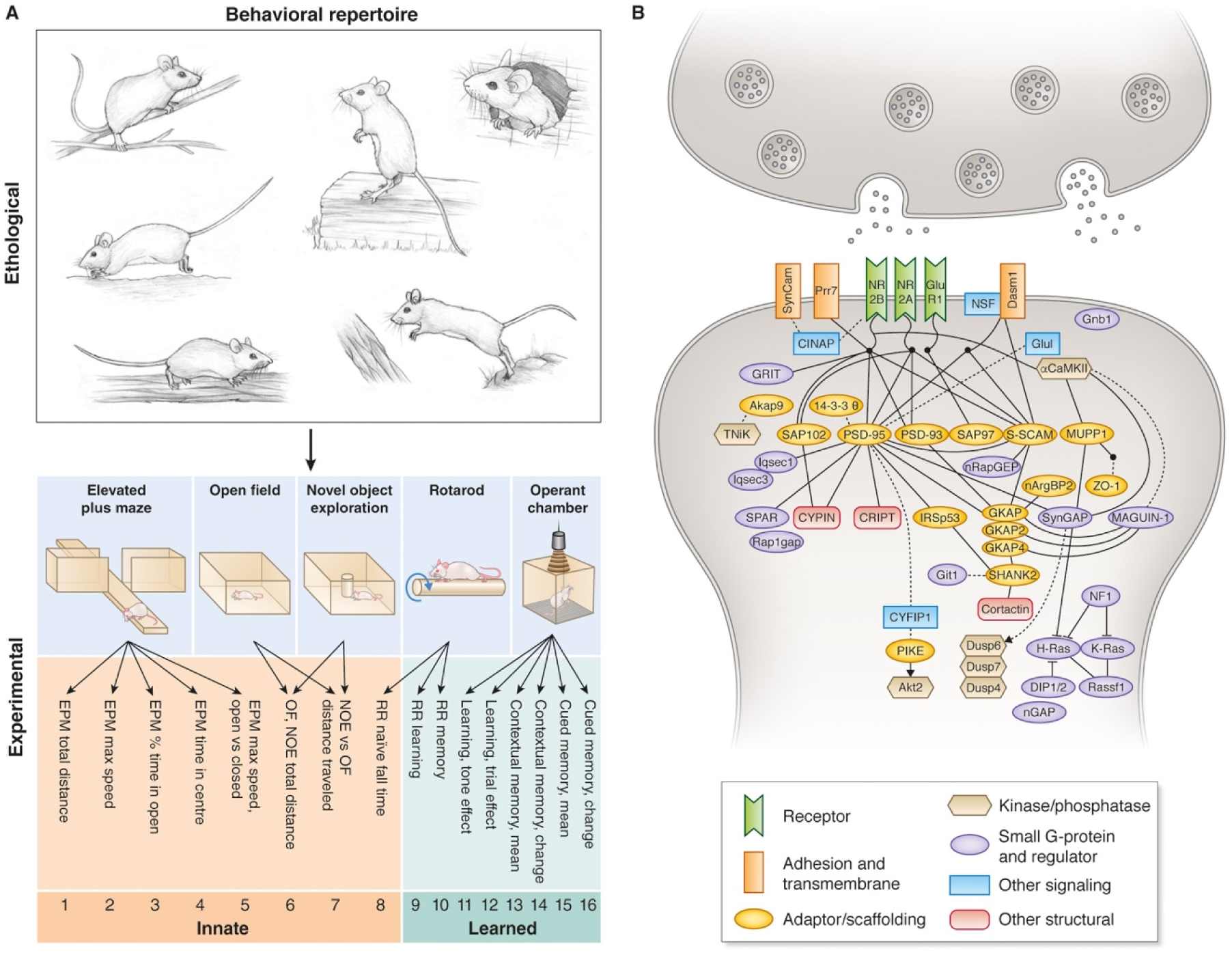
Postsynaptic genetic dissection of a repertoire of innate and learned behaviors. A. Ethologically relevant behavioral responses (top) were assessed with experimental apparatus (bottom) probing innate and learned responses. Sets of eight innate and eight learned responses comprised the overall repertoire. B. The postsynaptic proteins mutated in this study. Protein interactions are indicated (solid lines, binary interactions; dotted lines, other functional interactions). Key, major protein classes.

**Figure 3.**
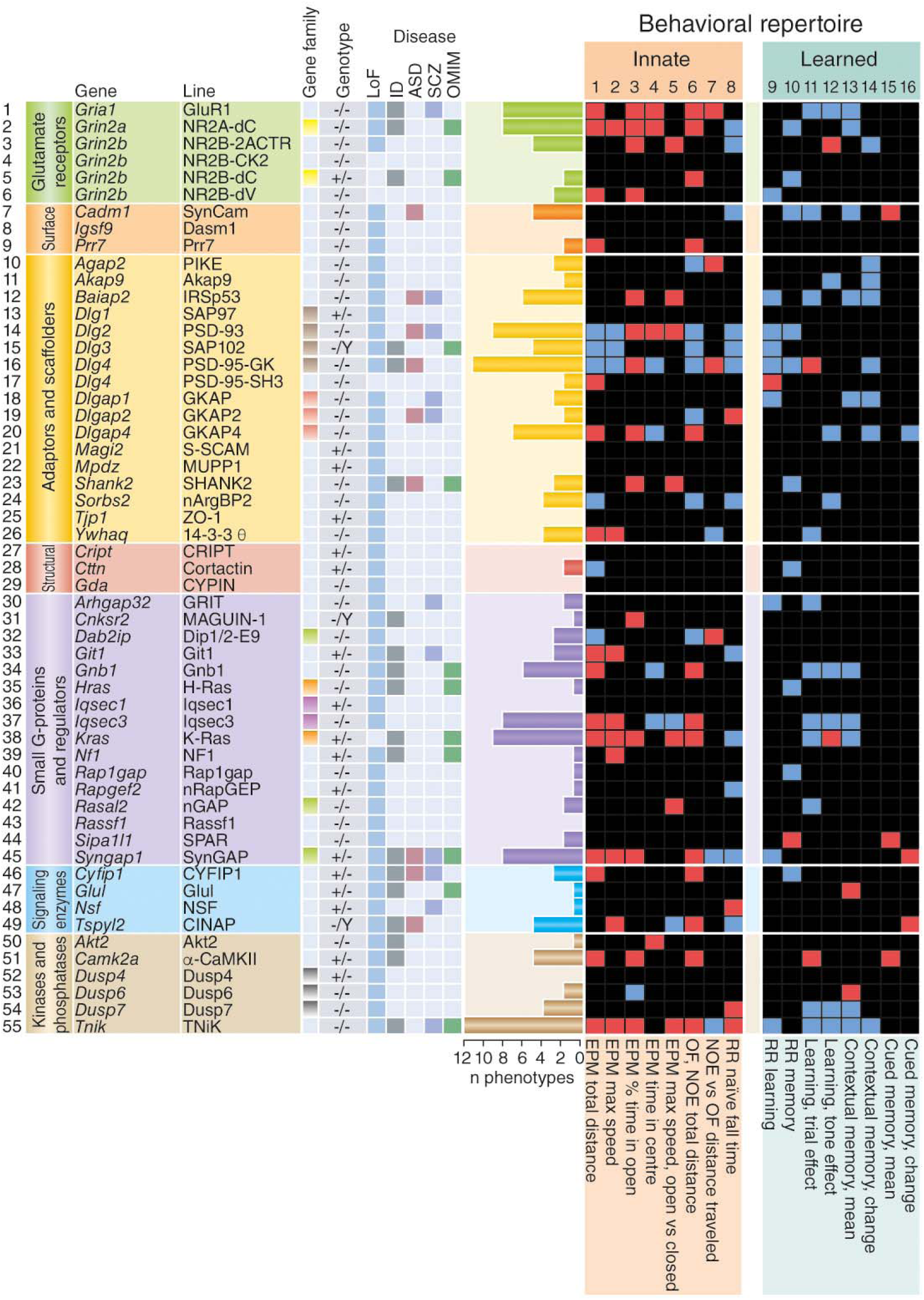
Summary of mutant phenotypes. Here, and in subsequent figures, phenotypes indicated by red and blue boxes illustrate mutation-induced amplification and attenuation of the parameters (*P* < 0.05), respectively. Black boxes indicate behaviors that were not changed significantly by mutation. Histograms show the total number of phenotypes for each mutant line. Innate (1–8) and learned (9–16) behavioral repertoire components are indicated. Gene family, identically colored boxes indicate paralogs from the same gene family; Genotype, homozygous, heterozygous or hemizygous for X-linked genes; LoF, loss-of-function alleles (50); Orthologs of human disease genes: ID, intellectual disability; ASD, autism spectrum disorder; SCZ, schizophrenia; OMIM, neural diseases from Online Mendelian Inheritance of Man; RR, rotarod; NOE, novel object exploration; OF, open field;EPM, elevated plus maze.

In total, 290,850 behavioral measurements were made from 2,770 mice. Cohorts of 10–20 mutant and 10–20 wild-type mice from each line (5–10 male and 5–10 female mice in each line, except for X-linked genes when only male mice were assessed) were examined in a battery of behavioral tests. The battery utilized the elevated plus maze (EPM), open field (OF), novel object exploration (NOE), rotarod (RR) and a chamber for cue and contextual fear conditioning (FC), which together present a diversity of elementary physical features and stimuli that elicit behavioral responses that are integral to more complex natural environments (Fig. 2A). From 105 automated behavioral measurements collected from each mouse, we identified a repertoire of 16 minimally correlated variables consisting of eight innate and eight learned responses (Fig. 2A, Table 1, Methods), as described and validated elsewhere^29^. This validation revealed that effect sizes of mutations were similar between C57BL/6J and 129S5 strains in our 16 behavioral variables. Hereafter, we refer to the 16 measures as the Overall Behavioral Repertoire (OBR), and this is subdivided into the Innate Behavioral Repertoire (IBR, components numbered 1–8 in figures) and the Learned Behavioral Repertoire (LBR, numbered 9–16), respectively. The significance of mutant/wild-type differences for each line was computed using two-factor (sex, genotype) ANOVAs. For each mouse, a phenotype score for each of the 16 behavioral components was obtained (Supplementary Table 2) and used to derive aggregate OBR, IBR and LBR scores^29^ (Methods).

**Table 1.** G2C experimental behavioral repertoire. EPM, elevated plus maze; OF, open field; NOE, novel object exposure; RR, accelerating rotarod; Learning consisted of three trials involving pairing of a mild foot shock with an audible tone; Contextual or cued memory ‘mean’ represents an average percentage freezing level during the indicated time period; Contextual or cued memory ‘change’ represents a difference in freezing between the first and last 30 s time bin of the time period indicated.

| Repertoire component |  | Variable name | Description |
| --- | --- | --- | --- |
| Innate | 1 | EPM total distance | Total distance (cm) travelled in EPM |
|  | 2 | EPM max speed | Maximum speed (cm/s) in EPM |
|  | 3 | EPM % time in open | % time in open vs. all arms of EPM |
|  | 4 | EPM time in centre | Time (s) in central zone of EPM |
|  | 5 | EPM max speed, open vs. closed | Difference in max speeds (cm/s) in open vs closed arms of EPM |
| | 6 | OF, NOE total distance | Total distance ($\log_{10}$ cm) in open field without and with novel object |
|  | 7 | NOE vs. OF distance travelled | Difference in distance travelled (cm) without and with novel object |
| | 8 | RR naïve fall time | Naïve fall time on rotarod ($\log_{10}$ s) |
| Learned | 9 | RR learning | Increase in fall time per trial (s/trial) in session 1 |
|  | 10 | RR memory | Excess fall time at mid-session 2 vs. mid-session 1 |
|  | 11 | Learning, trial effect | % time freezing before 3rd trial vs before first trial |
|  | 12 | Learning, tone effect | % time freezing attributable to third tone vs first tone |
|  | 13 | Contextual memory, mean | Mean % freezing during first 120 s re-exposure to context vs. first 120 s on previous day |
|  | 14 | Contextual memory, change | Increase in % freezing during 120 s re-exposure to context |
|  | 15 | Cued memory, mean | Mean % freezing due to 120 s tone re-exposure vs. initial tone on previous day |
|  | 16 | Cued memory, change | Increase in % freezing during 120 s re-exposure to tone |

### Combinations of postsynaptic proteins regulate repertoire components

The phenotypes of 55 lines of mice in each of the behaviors comprising the OBR, IBR and LBR are summarized in Figure 3. We were surprised to find that both the IBR and LBR were affected by mutations in genes encoding postsynaptic proteins because it is often posited that most postsynaptic signaling proteins are primarily involved with learning and memory. Furthermore, the mutations were not exerting pervasive effects as only 20.3% of all possible phenotypes (162/800, *P* < 0.05; 11.1% at *P* < 0.01) were observed in the 50 LoF mutations. Sexually dimorphic phenotypes (sex × genotype interaction) were rare, comprising 1.5% in 47 LoF autosomal lines (11/752 at *P* < 0.01; 6.3% at *P* < 0.05) (Supplementary Fig. 1, right-hand panel), and because this was close to the false positive rate, we restricted further analysis to main genotype effects.

The pattern of phenotypes revealed that each component of the repertoire required combinations of particular postsynaptic proteins (Fig. 3). Moreover, the combination regulating each behavior was subdivided into mutations that amplified (red boxes) or attenuated (blue boxes) the magnitude of the response. We next asked if the combinations were restricted to any particular class of protein and found that all seven classes contributed to both IBR and LBR, with individual behavioral components typically using the majority of the seven classes (Fig. 3). This indicates that each behavioral response requires the integrated role of combinations of all the major protein classes in the postsynaptic proteome.

Although each behavioral component required a unique combination of proteins, most proteins regulated more than one behavior. Indeed, the neurotransmitter receptor subunits of the NMDA (NR2A and NR2B) and AMPA (GluR1) receptors were required for both innate and learned behaviors, consistent with previous reports^30–32^ (Fig. 3). By contrast, eight proteins (Akt2, MAGUIN-1, H-Ras, NF-1, Rap1gap, nRapGEP, NSF and Glul) affected only a single repertoire component. Thus, each behavioral response is typically regulated by proteins that participate in the regulation of several other behaviors together with proteins that are more restricted to that particular behavior. To further examine the similarity in the molecular mechanisms of innate and learned behaviors, we counted IBR and LBR phenotypes and found that they were similarly affected: 98 IBR phenotypes compared with 64 LBR phenotypes (using the *P* < 0.05 cutoff) were observed (after correcting for false positives, odds ratio = 1.27, *P* = 0.15; chi-squared test) in the 50 LoF lines. These findings indicate that combinations of postsynaptic proteins from multiple classes regulate instinctive and learned responses.

We found that some behaviors were relatively frequently disrupted by mutations. In the IBR, locomotor activity was affected by mutations in 36% of the genes (total distance travelled: EPM, 18 phenotypes and OF/NOE, 18 phenotypes), motor ability was affected by 30% of genes (15 phenotypes in RR naïve fall time), and anxiety levels were regulated by mutations in 26% of genes (13 phenotypes in “EPM % time in open”). In the LBR, contextual fear memory was affected by mutations in 22% of genes (“Contextual memory, mean”, 11 phenotypes), motor learning was altered by mutations in 20% of genes (“RR learning”, 10 phenotypes), whereas cued fear memory was influenced by the fewest number of genes (“Cued memory, mean”, 6%, 3 phenotypes).

### A subset of postsynaptic proteins controls the behavioral repertoire

The identity of the postsynaptic proteins of greatest behavioral importance has remained unknown because previous studies lacked methodological standardization and rarely examined the roles of more than a few proteins. Quantitative comparison of phenotypes is also essential for understanding how proteins work together. Using our standardized dataset, we asked which proteins have the strongest effect on the OBR. The effect size (Cohen’s d) values of the 16 phenotypes for 50 LoF mutations were aggregated to generate an OBR score and ranked accordingly (Fig. 4A). PSD95 and TNiK mutants had the strongest phenotypes, implicating these two proteins as major regulators of many behaviors. Following these, in decreasing order of phenotypic severity, were mutations in AMPA receptor subunit GluR1, scaffold proteins (PSD93, SAP102, IRSp53) and signaling proteins (SynGAP, CINAP, Iqsec3, Gnb1) in the top ten. The next group of ten included mutations in NMDA receptor subunits (NR2A, NR2B), α-CaMKII, Shank2 and GKAP4, among others.

**Figure 4.**
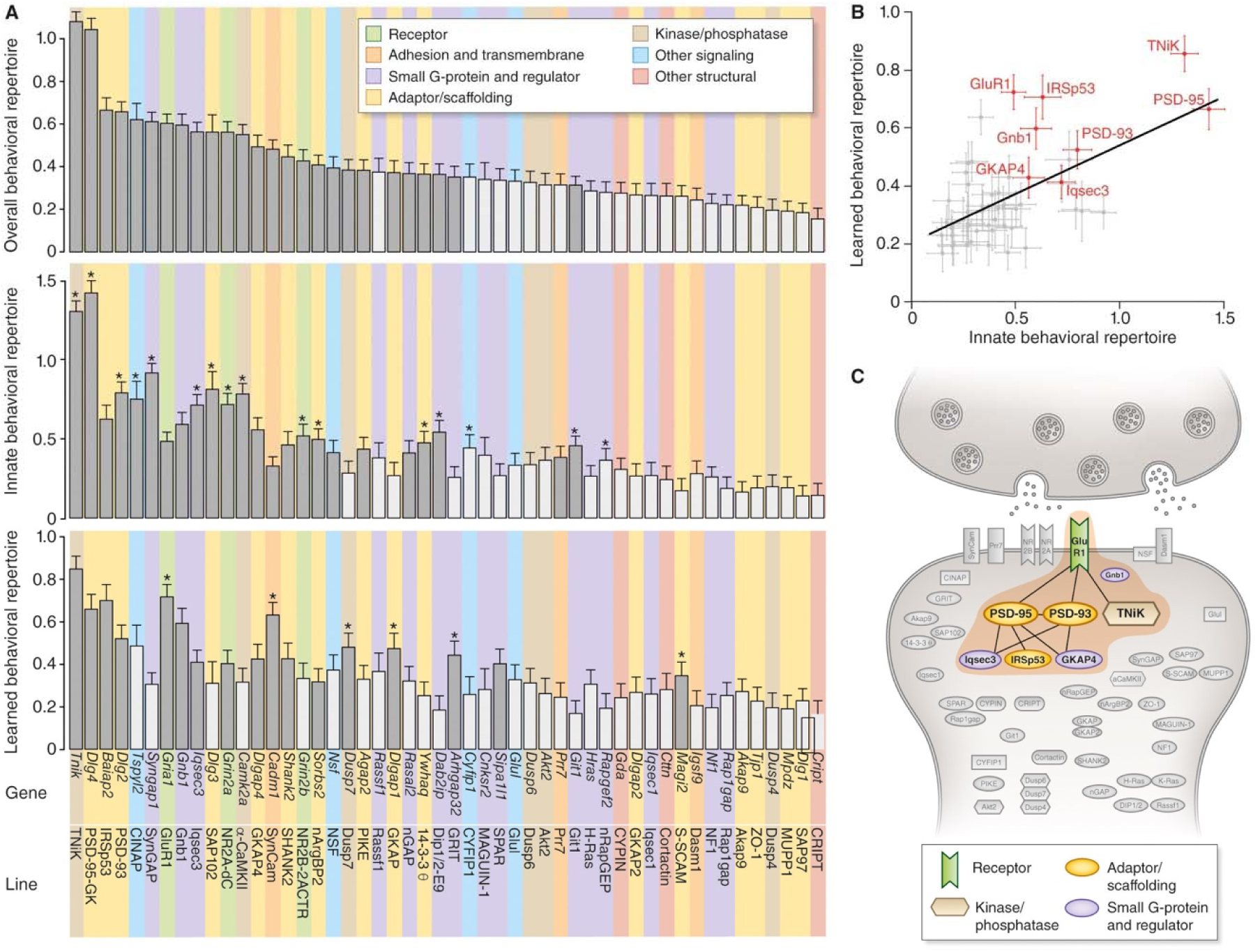
Major effect proteins in innate and learned behavior. A. (Top) Overall Behavioral Repertoire (OBR) phenotypes ranked by effect size (Cohen’s d) values for 50 LoF mutations. (Middle) Innate Behavioral Repertoire (IBR) phenotype effect size values. Asterisks indicate genes biased to affect innate phenotypes. (Bottom) Learned Behavioral Repertoire (LBR) phenotype effect size values. Asterisks indicate genes biased to affect learning and memory phenotypes. Shaded bars represent average effect sizes that are larger than expected by random chance (*P* < 0.05). Error bars represent standard error of the Cohen’s *d* estimate. B. Correlation between phenotype effect size (Cohen’s d) values of innate and learned behavioral repertoires. Spearman rho = 0.491, *P* = 0.00035. Eight proteins (red) were required for a significant correlation. Error bars represent standard error of the Cohen’s *d* estimate. C. The eight core proteins (brown zone) controlling the innate and learned behavioral repertoires, centered on PSD95, are surrounded by satellite proteins.

Next, we examined the IBR and LBR aggregate scores separately and found that some proteins were biased toward a role in either innate or learned behaviors (Fig. 4A). Sixteen mutations were significantly biased toward disrupting innate behaviors and six were biased to alter learned behaviors (*P* < 0.05, permutation tests; asterisks in Fig. 4A). For example, SynGAP, SAP102 and α-CaMKII showed substantially greater impacts on innate behaviors, whereas SynCam, Dusp7, GKAP1 and GRIT showed a greater impact on learned behaviors.

To identify those proteins that play the most important roles in both the IBR and LBR, we compared their behavioral scores for the 50 LoF lines (Fig. 4B). We found a significant correlation (Spearman rho = 0.491, *P* = 0.00035) and identified eight proteins (PSD95, TNiK, PSD93, GluR1, Gnb1, IRSp53, GKAP4, Iqsec3) of major effect that were essential for the correlation. We conclude that this combination of eight proteins is of major importance and forms the core of the shared molecular machinery of both innate and learned repertoires. The other proteins are more specific to individual behavioral components. We hereafter refer to the two sets as the “core” and “satellite” proteins, respectively.

Strikingly, 7 of the 8 proteins in the core set are known to be highly interconnected (Fig. 1B) and co-purify in the multiprotein complexes assembled with PSD95^13,19–21,33,34^ (Fig. 4C). These high molecular weight (~1.5 MDa) PSD95 supercomplexes have been isolated from the mouse and human brain and found to contain diverse functional classes of proteins including neurotransmitter receptors, ion channels, adhesion, signaling and structural proteins^13,19–21,33,34^. Although the eighth protein of the core set, Iqsec3, was not found in these studies, both Iqsec1 and Iqsec2, which are Iqsec3 paralogs, were reported. Thus, the genetic screen of behavioral phenotypes identified proteins in PSD95 supercomplexes as central components of the postsynaptic machinery that controls the behavioral repertoire.

The postsynaptic terminal contains other multiprotein complexes, such as the ~350 kDa complexes assembled by SAP102^19,34^. Our data show that SAP102 has a lower OBR score than PSD95, and that SAP102 primarily plays a role in the IBR (Fig. 4A). These data suggest that multiple postsynaptic complexes are building blocks of the components of the behavioral repertoire, with PSD95 supercomplexes playing the major role.

### Genetic tuning of behavioral optima

The magnitude of behavioral responses is crucial to the survival of an animal and the genetic and synaptic mechanisms establishing this key feature of behavior are poorly understood. Our quantitative phenotyping revealed that each behavioral component of the repertoire is attenuated or amplified by different postsynaptic mutations (Fig. 3, blue and red boxes respectively), indicating that these genes control a set-point, or optimum, of each behavioral response. To examine this in more detail, we counted LoF mutations that amplified or attenuated each behavior (Fig. 5A). Surprisingly, the set-points for most (7/8) of the innate behaviors were disrupted with similar frequency in both directions, whereas most (5/8) learned behaviors showed unidirectional biases. To further assess the bias in each variable, we asked how often phenotypes were in either the majority or minority direction (Figure 5A). For the IBR, 68 and 30 phenotypes (ratio of 2.3 to 1) were in the majority and minority directions, respectively, whereas for the LBR there were 57 and 7 (ratio of 8.1 to 1) phenotypes in the majority and minority directions. These results demonstrate strong bias toward unidirectional phenotypes in learning and memory variables (odds ratio 0.28, *P* = 0.0039, Fisher’s exact test), revealing that mutations more frequently lead to impairments (rather than improvements) in learning and memory.

**Figure 5.**
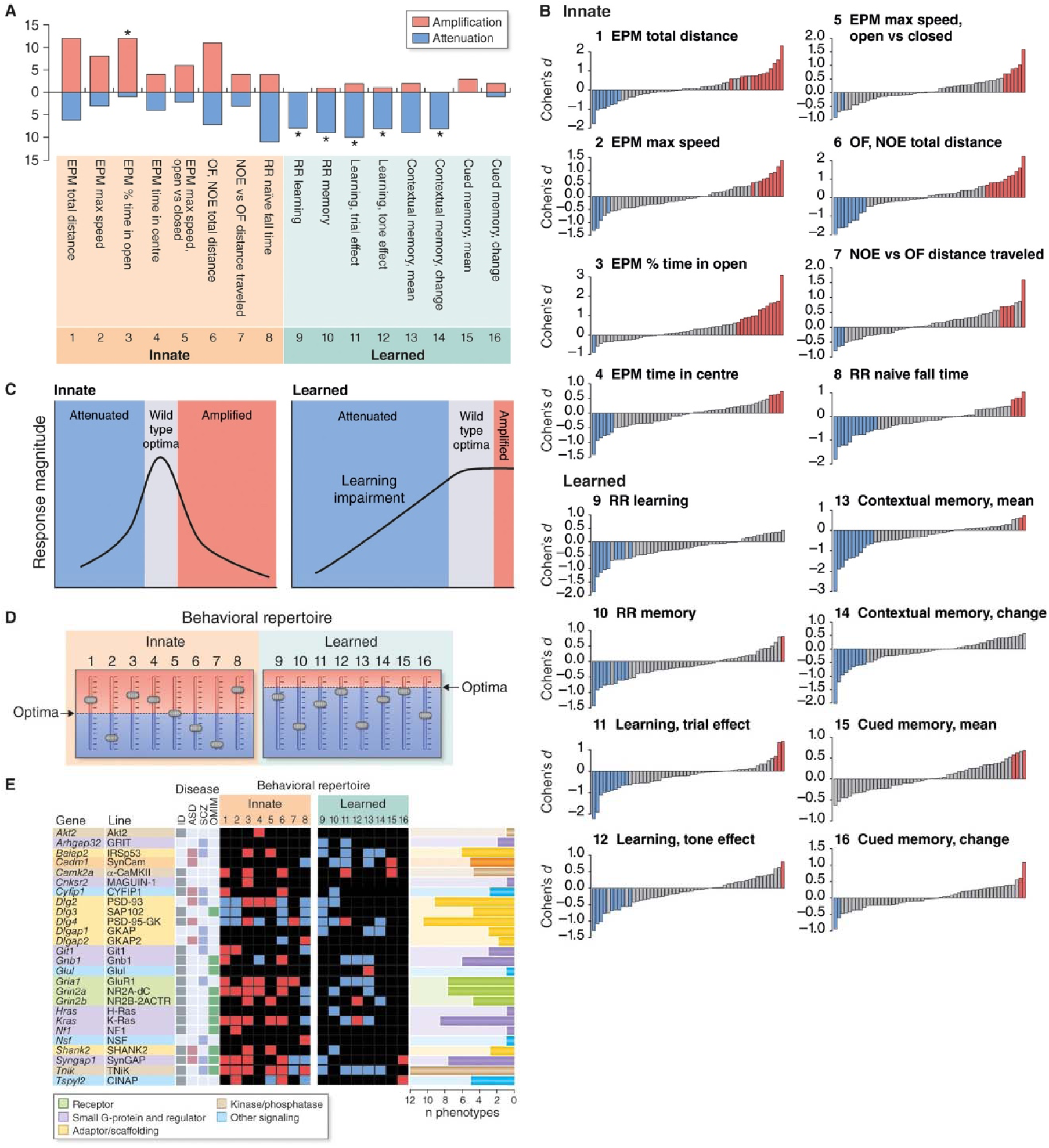
Tuning of response optima and disease. A. Number of LoF mutations amplifying or attenuating each behavior. Directional bias indicated as follows: *\*P* < 0.05, binomial test, assuming no bias. B. Ranking of LoF mutations by the direction and phenotype effect size (Cohen’s *d*) value of each behavioral variable. C. Genetic tuning of behavioral optima. Innate behavioral responses are tuned within a broad bidirectional range, whereas learned responses are at near-maximal optimum. Mutations shift the behaviors from the wild-type optima. D. Set-points and optima of the magnitude of each behavioral component. The position of sliders indicates the shift from the optima caused by a mutation. E. Behavioral repertoire phenotypes (*P* < 0.05) of intellectual disability (ID), autism spectrum disorder (ASD) and schizophrenia (SCZ) orthologs are indicated by red (amplified behavior response) or blue (attenuated) bars. Black bars, non-significant.

These findings indicate that the molecular mechanisms of the postsynaptic proteome have evolved to a near-maximal learning optimum, whereas the optima of innate components are set within a broad and bidirectional range (Fig. 5C). This observation is reinforced by rankings of the bidirectional phenotype effect sizes for each behavioral measurement (Fig. 5B, Supplementary Figs. 2 and 3). This means that mutations will typically cause learning impairments, whereas innate responses can be either amplified or attenuated. For example, mutations in PSD95 or TNiK both result in attenuated contextual memory but have opposite locomotor phenotypes (hypo- and hyperactivity, respectively, in OF/NOE; Fig. 3). There was only one innate behavior that showed a unidirectional phenotype (bias *P* = 0.0034, chi-squared test): an anxiety measure (EPM % time in open), which was reduced in 12 mutants and increased in only one. This suggests that learning disability and reduced anxiety are typical phenotypes and occur alongside other innate phenotypes. Figure 5D schematically illustrates the concept of optimal set-points for innate and learned behaviors.

### Disease mutations restructure the behavioral repertoire

Mutations and genetic variation in postsynaptic proteins cause more than 130 monogenic and polygenic brain disorders, including schizophrenia, depression, autism and intellectual disability^11,20–25^. Our datasets included mutations in 26 orthologs of intellectual disability (ID), autism spectrum disorder (ASD) and schizophrenia (SCZ) susceptibility genes, and we have compared the innate and learned behavioral phenotypes in these mutant mice (Fig. 5E, Supplementary Table 3). It is important to note that this is the first study that makes a systematic and standardized comparison of many gene mutations linked to psychiatric disorders. As shown in Figure 5E, many of the innate and learned components of the behavioral repertoire were shifted from their optima in these mouse mutants. Learning impairments and increased or reduced anxiety were common phenotypes. There was no clear pattern of phenotypes associated with the classifications of any one of the classes of disease genes. Together, these findings indicate that these diseases are associated with a reconfiguration of multiple components of the behavioral repertoire, with different combinations of behaviors having aberrant or maladaptive responses.

### Evolution of the behavioral repertoire

Understanding how the sophisticated behavioral repertoire of humans and other vertebrates evolved is a major scientific question. There have been two landmarks in the molecular evolution of the postsynaptic proteome. The first is the origin of the protein classes, which arose in unicellular organisms several billion years ago^35^. The second was the expansion in the complexity of the postsynaptic proteome early in the vertebrate lineage ~550 million years ago, secondary to two whole-genome duplications that generated combinations of paralogs (ohnologs) in each protein class^36,37^. Our large-scale data allowed us to gain insight into how each of these molecular evolutionary events impacted the behavioral repertoire of extant mammals.

We first compared the behavioral repertoire of paralogs from six different classes of postsynaptic proteins (Fig. 6A). Within each gene family, mutations in paralogs showed distinct phenotypes, indicating that specialized non-redundant functions arose in vertebrates after the genome duplication events. Thus, the vertebrate genome duplications and consequent expansion in postsynaptic proteome complexity has contributed to more complex molecular regulation of the set-points of behavioral responses. Next, we compared the repertoire of homologs that arose in vertebrates, invertebrates and unicellular eukaryotes (Fig. 6B), revealing that all these proteins play a role in innate and learned behaviors. Thus, ancient and recent molecular signaling mechanisms contribute to the control of the behavioral repertoire of the mouse.

**Figure 6.**
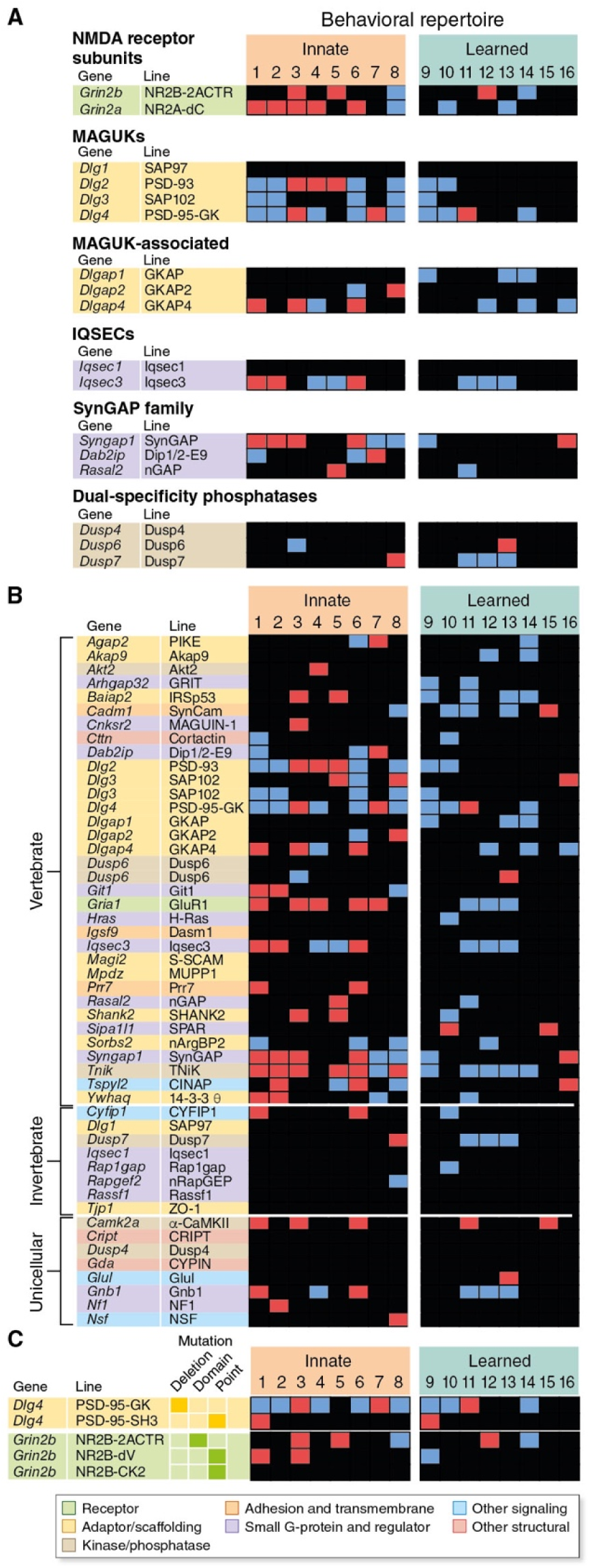
Genome evolution shaped the behavioral repertoire. A. Behavioral repertoire phenotypes caused by mutations of vertebrate paralogs in six protein classes. B. Behavioral repertoire phenotypes plotted separately for mutations in homologs of unicellular eukaryote, invertebrate and vertebrate genes. C. Behavioral repertoire phenotypes caused by different structural mutations (gene deletion, domain deletion or point mutation) in PSD95 and NR2B.

The genomic mechanisms for paralog diversification include single nucleotide polymorphisms (SNPs) that can induce fine structural changes in proteins^38^. Postsynaptic proteins were highly constrained during vertebrate evolution^11^, so even small structural changes by SNPs may cause deleterious phenotypes. It was therefore of interest to contrast the impact on the behavioral repertoire of fine structural changes with that of whole gene deletions. We first compared the deletion of PSD95 with a SNP that interferes with the function of the SH3 domain^39^ (Fig. 6C): the point mutation affected two behaviors in contrast to the deletion that affected 11 behaviors. Next, we examined three alleles of *Grin2b* that modified the cytoplasmic domain by a complete substitution^40^, a small deletion and an SNP (Supplementary Data 1), respectively: the less severe mutations showed fewer phenotypes (Fig. 6C). These results demonstrate that small structural changes in key postsynaptic proteins can fine-tune specific behavioral responses.

## Discussion

### A molecular mechanism for a versatile behavioral repertoire

We have conducted a large-scale genetic analysis that explores the role of the postsynaptic proteome in a set of innate and learned responses to environmental stimuli. Remarkably, a unique combination of proteins was required for each innate or learned response. The postsynaptic proteome contains over 1,000 proteins^10–17^ and thus, the combinatorial possibilities could potentially generate a vast behavioral repertoire. We found the behavioral diversity was increased by the expansion in the complexity of the postsynaptic proteome that arose from ancient genome duplications. These findings provide robust evidence that the complexity of the postsynaptic proteome generates a rich and diverse behavioral repertoire in vertebrate species.

Understanding the relationship between innate and learned behavior has been a central question in psychology, and Lorenz argued that this dichotomy could only be understood by the discovery of the mechanisms underlying these two classes of behavior^7^. Our findings demonstrate that innate and learned behavior share a common synaptic mechanism in the combinatorial use of the same sets of postsynaptic proteins and PSD95 supercomplexes. It is the protein combination rather than the individual proteins that distinguished innate from learned behavior. This is illustrated by SynGAP, SAP102 and α-CaMKII, which have been previously shown to play a role in learning^41-43^, but quite clearly play a role in innate behaviors too. Indeed, in our behavioral repertoire these proteins had a greater role in innate behaviors. By contrast, proteins such as SynCam, Dusp7, GKAP1 and GRIT were biased to learned behaviors.

The combinatorial molecular mechanism also explains how the magnitude of behavioral responses is genetically controlled, something crucial for survival and adaptation to environmental niches. For each behavioral response, we found that the combinations of proteins were divided into those that enhanced or attenuated the response magnitude. This suggests that the concerted action of these proteins ‘tunes’ the response magnitude. Thus, the combinatorial mechanism is highly versatile in that it can regulate many individual behaviors as well as their response magnitude.

### Postsynaptic complexes and synaptome organization

The most severe behavioral perturbations were caused by proteins encoding components of PSD95 supercomplexes. These high molecular weight multiprotein complexes encompass neurotransmitter receptors, ion channels and signaling proteins^13,19–21,44,45^ that control short- and long-term synaptic plasticity^28,42,46,47^. Important components of these supercomplexes include the NMDA receptor and proteins interacting with AMPA receptors^13,19,21^. Our results revealed that different behaviors used different combinations of PSD95 supercomplex proteins, suggesting that subtypes of supercomplexes^44^ composed of different proteins selectively participate in specific behaviors. In addition to the “core” role of PSD95 supercomplexes in the behavioral repertoire, we found that each behavior used “satellite” proteins found in other postsynaptic complexes, such as the ~350 kDa complexes organized by SAP102^19^. Thus, PSD95 supercomplexes and other postsynaptic signaling complexes are structural components of the postsynaptic proteome that together orchestrate the behavioral repertoire.

Synaptome mapping of the molecular composition of individual synapses across the whole mouse brain reveals that PSD95 and SAP102 are differentially distributed into diverse synapses containing both or either protein^48^. Furthermore, each region of the brain shows a characteristic combinatorial postsynaptic proteome expression profile^49,50^. Thus, the protein combinations revealed in our genetic study of behavior may reflect the distribution of different complexes into the molecular subtypes of synapses that populate the circuits mediating these behaviors.

### Diseases of the behavioral repertoire

Clinical studies show that psychiatric, neurological and developmental disorders arising from mutations and genetic variation typically present with a constellation of behavioral phenotypes^51^. There are five findings from our study that provide a framework for understanding how diseases with a genetic etiology result in the pattern of behavioral disorders that characterize these diseases. First, most single gene mutations resulted in changes in multiple innate and learned behaviors. Thus, a disease mutation can be considered to restructure the behavioral repertoire. Second, each behavior was regulated by a combination of genes. This is consistent with the polygenic nature of human behavioral disorders that are known to target excitatory synapses^11,20–25^. Third, each mutation resulted in a unique overall change to the behavioral repertoire and many mutations had shared individual phenotypes. This is relevant to the clinical observation that these diseases manifest with a ‘spectrum’ of overlapping phenotypes. Fourth, mutations shifted the magnitude of innate and learned responses, consistent with the observation that human behavioral disorders are typically quantitative or graded changes of normal behaviors. For example, anxiety is a common and disabling human condition and we observed that over 25% of the mutations in our study resulted in anxiety phenotypes of different magnitudes. Furthermore, most postsynaptic mutations result in learning impairment, a common human phenotype. Fifth, our data show that subtle structural mutations result in fewer changes to the behavioral repertoire than larger structural mutations. This may be relevant to the smaller effect sizes of SNPs in common polygenic disorders.

Our datasets and phenotyping methods can be applied to studies of human genetic diseases with a view to potential therapies. For example, it is possible to ask if the combinations of genes that define each behavior in mice can be associated with specific human phenotypes or disorders. Many studies using mouse models of human disease have focused on a particular behavioral test and our results reveal that behavioral phenotypes affecting many components of the behavioral repertoire typically occur. Thus, a test battery approach will be less likely to miss phenotypes and provide a more comprehensive analysis of the disease model. This could be important in the context of therapeutic rescue experiments because a therapy that repairs every maladaptive behavior is more likely to be useful than one that rescues only a single behavior.

In conclusion, this study reveals genetic and synaptic mechanisms that can explain how the diversity and magnitude of innate and learned behaviors are individually controlled and coordinated into a behavioral repertoire. The combinatorial principles revealed by our analysis of a subset of the mouse behavioral repertoire are likely to apply to a much broader range of behaviors. This powerful mechanism provides a simple means by which mutations and variants arising in the genome shape the behavioral repertoire of vertebrate species.

In a companion manuscript we study the electrophysiological properties of synapses in the CA1 region of the hippocampus in the same set of mutant mice. This shows that the combinatorial mechanism defined here is important in computing the information encoded in patterns of neuronal activity and in controlling both short- and long-term plasticity^28^. We go on to integrate the behavioral and electrophysiological datasets to show that this species-conserved molecular mechanism converts the temporally encoded information in nerve impulses into the repertoire of innate and learned behavior.

## METHODS

### Animal models

Cohorts of homozygous and wild-type mice were generated from intercrosses of heterozygous animals. If homozygous mice were lethal or produced too few offspring, heterozygous mice were used. Within a line, mutant and wild-type mouse cohorts were litter-matched. Target group sizes for autosomal mutations were 20 mutants and 20 wild-type mice, with approximately 10 females and 10 males in each group. X-chromosome mutations were assayed in males only with group sizes of at least 10 wild-type and 10 mutant mice. For phenotypes of true size (Cohen’s *d* = 1 at α = 0.05), group sizes of 20 mutants and 20 wild-types confer 87.5% power to detect a phenotype, whereas 10 mutants and 10 wild-types confer 53.0% power. Details of knockouts, vector designs, genotyping, and breeding can be found in Supplementary Data 1 and online (http://www.genes2cognition.org/publications/g2c).

### Behavioral tests, protocols, and definition of a behavioral repertoire

Behavioral testing was carried out over five days in five standardized artificial environments. Details and validation of the protocol, as well as handling of the data, are described elsewhere ^29^. These environments addressed innate and learned responses to the environment, including sensorimotor function, perception, anxiety, novelty-seeking, locomotion, motor coordination and fear. A total of 105 highly redundant measures were made on each mouse using video and other tracking devices linked to specialized software. Of these, sixteen highly differentiated variables, which effectively summarized the 105 variables, were chosen and validated^29^.

### Elevated plus maze (EPM)

The elevated plus maze was the first test in the behavioral protocol. The apparatus was a plus-shaped maze with infrared illumination available from Tracksys (Nottingham, UK) with two exposed arms 45 cm above the ground and two arms protected with walls. Five variables were selected for analysis in the EPM. 1) “EPM total distance” is the total distance (cm) travelled in any arm or central zone of the EPM. 2) “EPM max speed” is the maximum speed (cm/s) travelled in any arm or central zone of the EPM. Although the maximum speed is expected to be correlated with the distance travelled, we used this variable to assess whether greater distance travelled was due to greater speed or because of generalized hyperactivity. 3) “EPM % time in open” is the percentage of time in the open arm compared with the time in any arm (exclusive of the central zone). This variable is a classical measure of the lack of anxiety. 4) “EPM time in center” is the total time (s) spent in the central zone of the EPM. 5) “EPM max speed, open vs closed” is the difference between the maximum speed (cm/s) observed in the open arms and the closed arms of the EPM. Speeds in the open arm are generally lower, presumably reflecting care taken by the animal in walking on the open arm or pauses to investigate the space around the open arm. This variable was used to assess additional aspects of the anxiety response in the open arm of the EPM.

### Open field (OF)

The open field test was conducted in the morning of the second day of the behavioral protocol. Animals explored the box for five minutes.

### Novel object exploration (NOE)

The novel object exploration task was carried out in the afternoon of the second day of the protocol. The same open field apparatus was used, but included an unopened aluminum 355 mL soft drink can placed in the center of the box. The animal explored the box for five minutes, and its behavior was recorded. Results from this assay and the open field were combined. Locomotion was measured as the total distance travelled during the open field and novel object exploration assays and denoted “OF, NOE total distance”. To measure mouse reaction to the change in environment, we chose the difference between the distance travelled in the novel object exploration and distance travelled in the open field, denoted “NOE vs OF distance travelled”.

### Rotarod (RR)

Animals were tested on the accelerating rotarod. Timing of a mouse’s fall was monitored by computer; latency to fall and maximum spindle speed were recorded for each trial. Each mouse underwent eight trials in the morning and eight in the afternoon. After each mouse, the apparatus was wiped down with ethanol wipes. Three behavioral variables describing each mouse’s rotarod performance were derived using linear models, and summarized all the trials of that mouse^29^. Briefly, “RR naive fall time” described innate motor coordination; “RR learning” measured the average number of extra seconds the animal stayed on the rotarod per training trial in the morning; and “RR memory” measured the difference between the mid-session fall time in the afternoon versus that in the morning. In this model of motor coordination, learning in wild-type mice is not correlated with naive performance, and memory is positively correlated with the measure of learning and more modestly with the measure of naive performance.

### Classical (fear) conditioning

Training was conducted on day 4 of the protocol. After two minutes of habituation, a 300 Hz tone at 83–86 dB was played for 30 s, co-terminating with a 2-s scrambled shock in the grid floor at 0.45 mA under control of Acctimetrics FreezeFrame software (Coulbourn Instruments, Whitehall, PA, USA). Two more tone-shock pairings were presented at 100 s intervals. The mouse’s behavior was recorded by an overhead video camera and freezing behavior was detected by Acctimetrics FreezeView software, version 2 (Coulbourn Instruments, Whitehall, PA, USA). Freezing behavior was recorded in 30 s time bins, except excluding 2 s of foot-shock during tone presentation. Testing was performed on day 5 in the same boxes; mice were placed in the box for three minutes to assess freezing in response to context, and then a tone was played for two minutes to assess cued freezing. Freezing was recorded in 30 s time bins. Six variables related to learning and memory were measured, as described^29^. “Learning, trial effect” measured the difference in freezing before the third training trial versus freezing before the first trial. “Learning, tone effect” measured the freezing response to the third tone versus the freezing response to the first. “Contextual memory, mean” measured the difference in mean freezing during the first two minutes of testing on day 5 versus freezing during the first two minutes on day 4. “Contextual memory, change” measured the increase in freezing from the first 30s time bin to the fourth, again comparing day 5 with day 4. “Cued memory, mean” measured excess freezing due to the tone on day 5, compared with excess freezing due to the first tone on day 4. “Cued memory, change” simply measured the change in freezing during the tone on day 5, comparing the first and last 30 s time bins during the tone. Expected linear relationships between learning variables (and separately between memory variables) were subtracted using linear models as described^29^. As a result, “Learning, trial effect” was weakly correlated to “Contextual memory, mean” and “Learning, tone effect” was weakly correlated to “Cued memory, mean”. In other words, two separable aspects of learning separately predict two aspects of memory. A caveat, however, is that these statistical manipulations are prone to over-fitting artefacts when calculated on smaller, commonly used cohort sizes.

### Use of Cohen’s *d* effect size to quantify effects of mutations on single variables and combinations of variables

Magnitude of the main genotype effect for each phenotypic measurement was assessed using Cohen’s d, a measure of standardized mean difference between groups (that is, the differences between the means divided by their pooled standard deviation) ^52^. Standard error of Cohen’s *d* for a phenotypic measurement was computed by sampling 1000 replicates from wild-type mice matched for genetic background, sex and group size. This sampling resulted in a set of 1000 Cohen’s *d* values distributed about zero. From this, the standard error of Cohen’s *d* was calculated as half of the spread between the 16^th^ and 84^th^ percentiles of this distribution (corresponding to the standard deviation in a standard normal distribution). Simulation of this procedure shows that these measures of standard error closely recapitulated those estimated by the accepted formula for Cohen’s *d* standard error. This strategy makes no assumption about the distribution of the underlying data. To assess overall impact of a mutation, Cohen’s *d* effect size magnitudes for multiple behavioral variables were averaged; this was done separately for eight innate and eight learned variables and the overall set of 16 behavioral variables. Again, standard error was computed by sampling wild-type animals. Due to sampling variation, this combined effect size is always positive, even when the true genotype effect sizes for component variables are zero. Therefore, significance of combined effect sizes was assessed in two steps. First, the null distribution of the combined phenotype effect size was computed by sampling from wild-type mice and the median value was identified. Significance of the combined effect size was computed from the number of standard errors by which it exceeded this baseline^29^.

## Acknowledgements

Mice from R. Sprengel and P. Seeburg (Gria1/GluR1, Grin2a/NR2A-dC, Grin2b/NR2B-dC), D. Bredt (dlg2/PSD93) and L. van der Weyden (Cadm1/SynCam, Rassf1/Rassf1). Computational support, J. Menendez Montes, J. Piatkowski, O. Sorokina; Technical support, V.J. Robinson, S. Syed Salim, G. Berry, E. Stebbings, E. Trenchard, R. Uren, H. Cooke, M. Green; Statistical advice, Ian J. Deary; Administration, J.V. Turner; Artwork, D. Maizels; Comments on manuscript, M. van Rossum, T.J. O’Dell, E. Fransén, M.V. Kopanitsa. Funding, Wellcome Trust, Wellcome Trust Sanger Institute and European Union Seventh Framework Programme under grant agreement numbers HEALTH-F2-2009-241498 (“EUROSPIN” project), HEALTH-F2-2009-242167 (“SynSys project”) and HEALTH-F2-2009-241995 (“GENCODYS project”).

## Author contributions

Gene targeting/mouse genetics, NHK, DJS, KES, DGF, EJT, KAE, TJR; Behavior experiments, LES, CMP, TJR, JN, LNL; Project tracking and colony management, KAE, MDRC; Statistical analysis, LNL; Transcriptome analysis, NGS; Database and website, MDRC; Direction and writing, SGNG.

## Data availability

The primary data and derived analyses are archived and available at http://www.genes2cognition.org/publications/g2c).

## Supplementary Materials

Fig S1-S3

Table S1-3

Supplementary Data File 1

