## Supplementary Figures and Tables for "Synaptic combinatorial molecular mechanisms generate repertoires of innate and learned behavior"

**Supplementary Figures and Table1**

### Supplementary Figure 1. Genotype and Sex × Genotype effect

Phenotypes in 55 lines of mice showing Genotype and Sex × Genotype effect. Note low frequency of sex × genotype effects. Behavioral phenotypes (P < 0.05) are indicated by blue (attenuated response) or red (amplified response) squares; black indicates no significant phenotype compared to wild-type mice. Behavioral repertoire components are: Innate (1, EPM total distance; 2, EPM max speed; 3, EPM % time in open; 4, EPM time in centre; 5, EPM max speed, open vs closed; 6, OF, NOE total distance; 7, NOE vs OF distance travelled; 8, RR naive fall time) and Learned (1, RR learning; 2, RR memory; 3, Learning, trial effect; 4, Learning, tone effect; 5, Contextual memory, mean; 6, Contextual memory, change; 7, Cued memory, mean; 8, Cued memory, change). The horizontal histogram shows the number of phenotypes for each gene/line. Columns: Protein class; Gene; Line, (mouse line name); Gene family (boxes of the same color indicate paralogs); Genotype (mutant genotype); LoF (loss-of-function alleles); Orthologs of human disease genes: ID, intellectual disability; ASD, autism spectrum disorder; SCZ, schizophrenia; OMIM, Neural diseases from Online Mendelian Inheritance of Man.

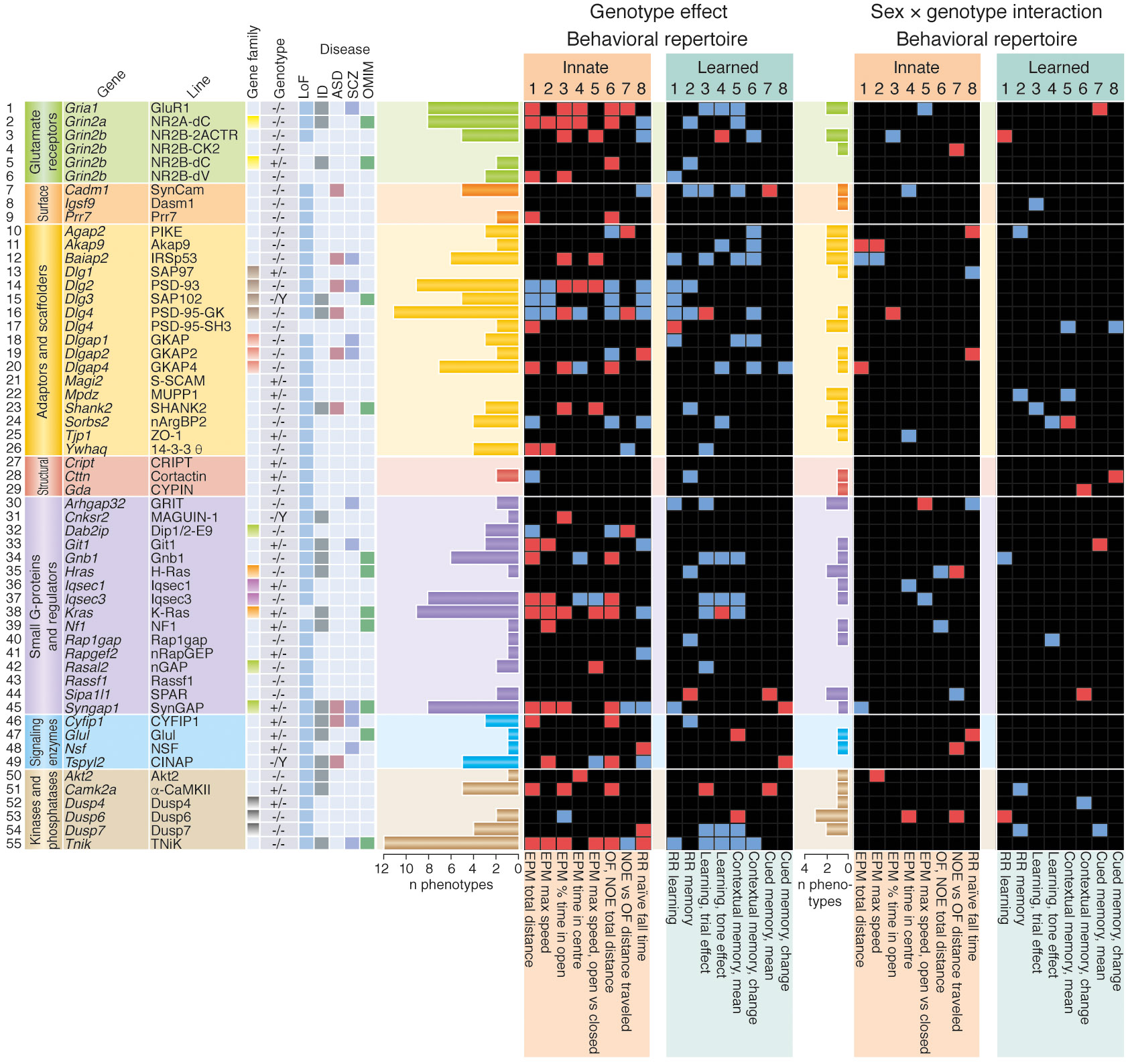

### Supplementary Figure 2. Innate behaviors

Ranked effect sizes (Cohen’s *d* values) and directions of eight innate behaviors in mice carying LoF mutations. Red and blue bars represent increased and decreased scores, respectively (*P* < 0.05), due to genotype effect determined by the two-factor (sex, genotype) ANOVA. Error bars represent standard error of Cohen’s *d* value (see Methods). Gene and Mouse line descriptions are as in Figure 2.

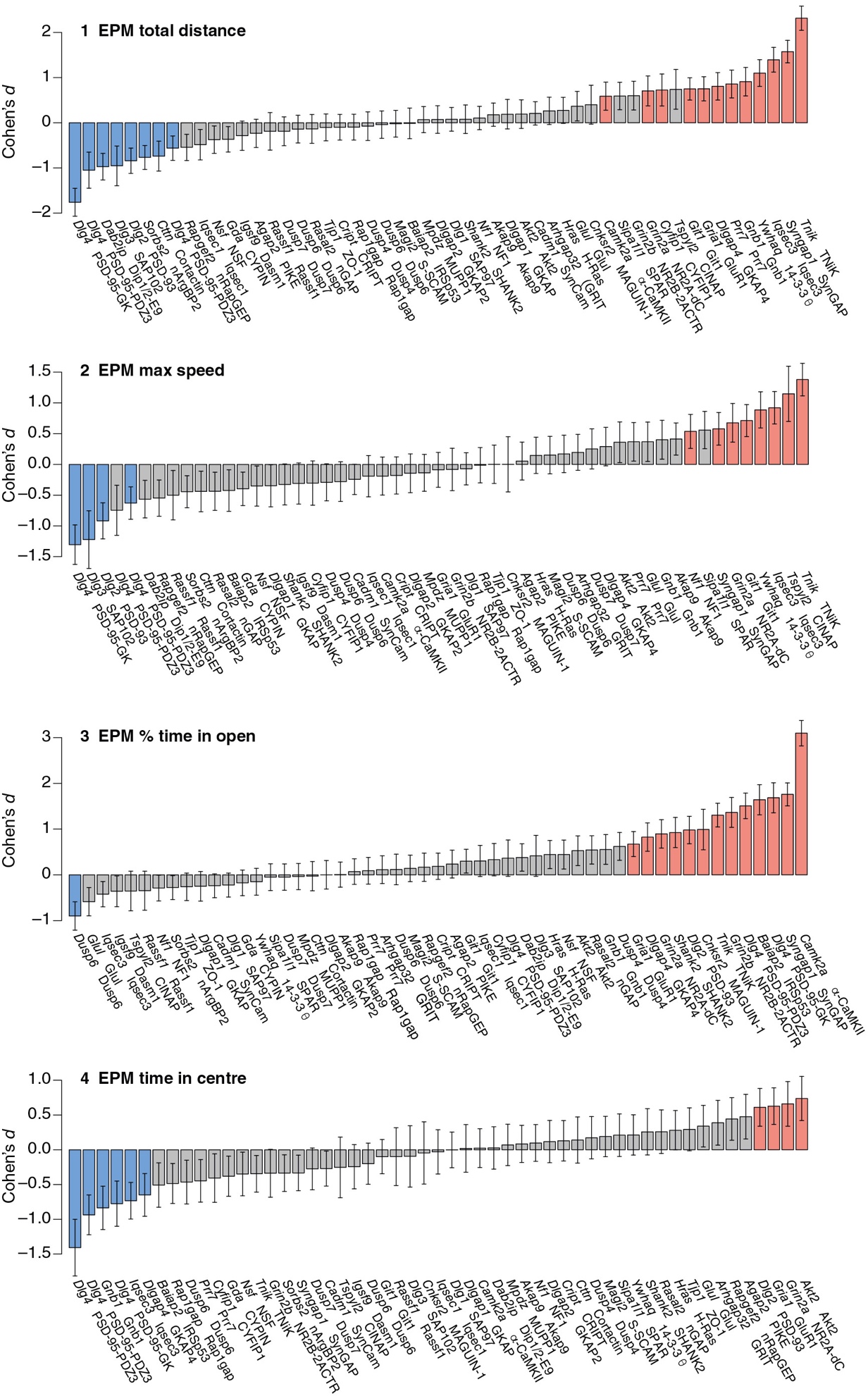

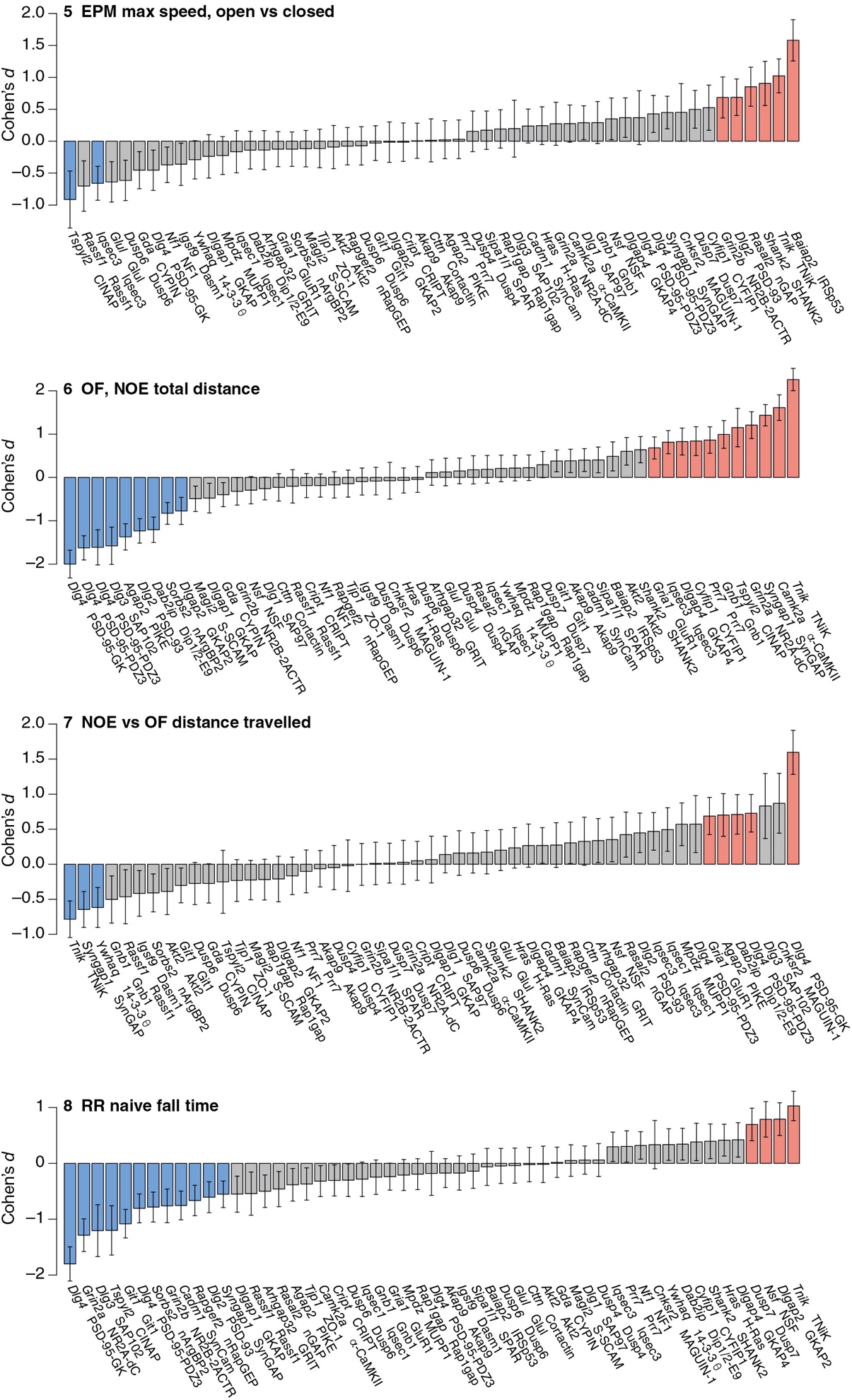

### Supplementary Figure 3. Learned behaviors

Ranked effect sizes (Cohen’s *d* values) and directions of eight learned behaviors in mice carying LoF mutations. Red and blue bars represent increased and decreased scores, respectively (*P* < 0.05), due to genotype effect determined by the two-factor (sex, genotype) ANOVA. Error bars represent standard error of Cohen’s *d* value (see Methods). Gene and Mouse line descriptions are as in Figure 2.

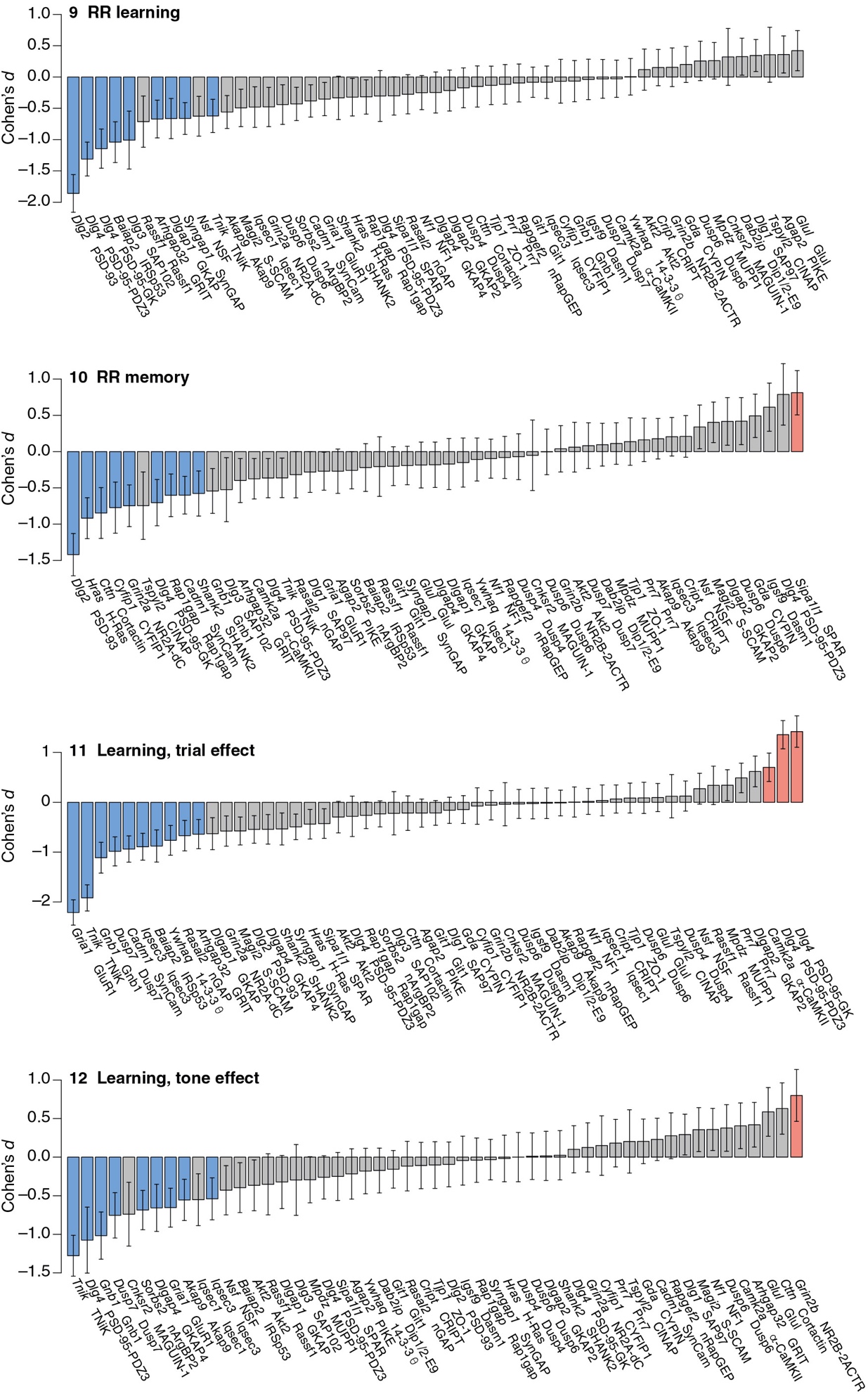

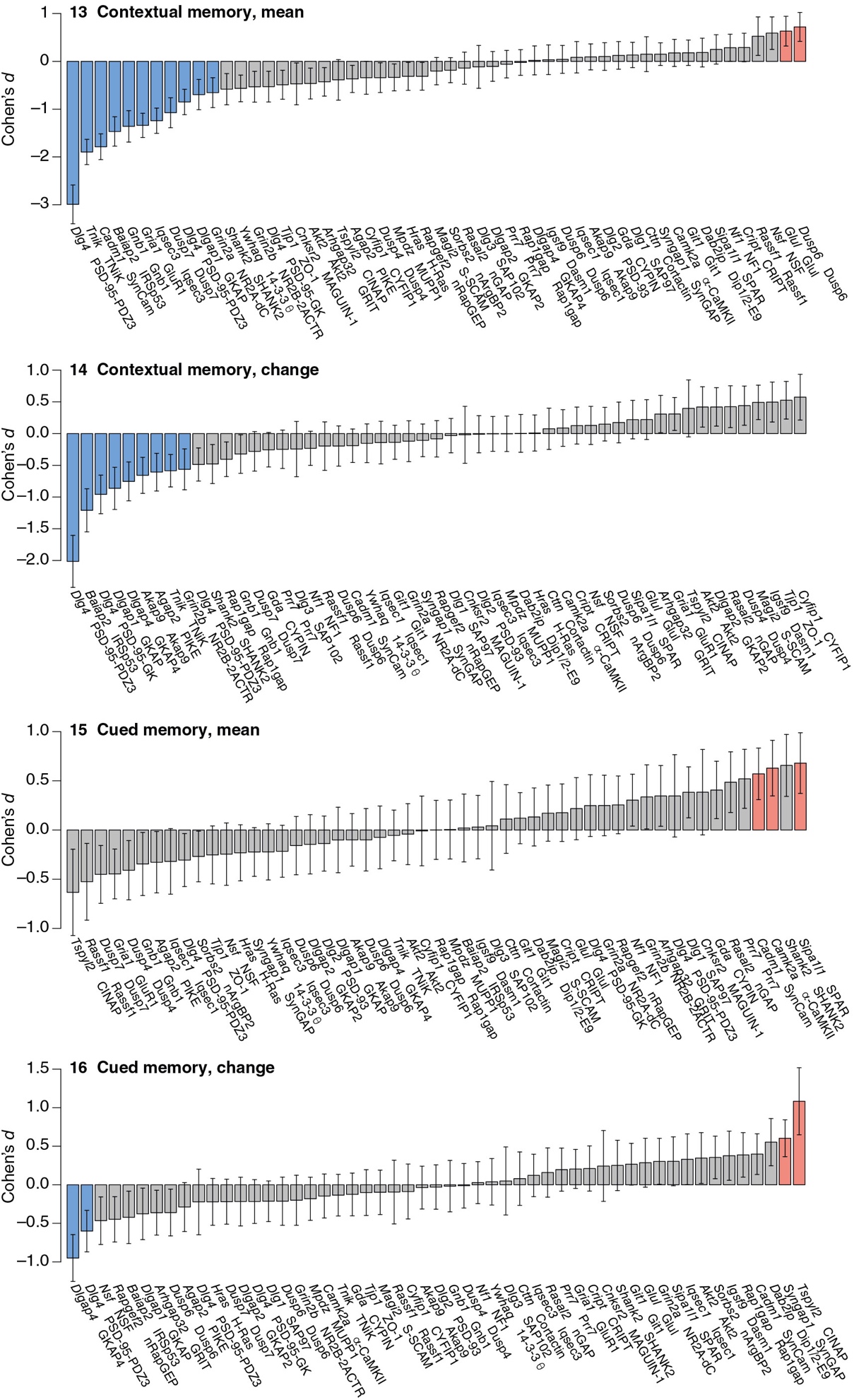

### Supplementary Table 1. Comparison of G2C phenotypes with published data

Comparison of behavioral phenotypes in the MGI database with G2C results shows 25/33 reported phenotypes agreed with phenotypes reported here. (see Supplementary Table 1.xls).

### Supplementary Table 2. Phenotype score for each of the 16 behavioral components.

Cohens *d* values for the 16 behavioural variables in each of the mouse lines studied (See Supplementary Table 2.xls)

### Supplementary Table 3. Human disease gene annotations

Disease gene annotations for the genes/mouse lines studied. References for intellectual disability (ID), autism spectrum disorder (ASD), schizophrenia (SCZ) shown and Online Mendelian Inheritance of Man (OMIM) reference numbers for neural diseases shown.

| **Gene** | **Line** | **ID** | **ASD** | **SCZ** | **OMIM neural disease** |
| --- | --- | --- | --- | --- | --- |
| *Akt2* | Akt2 | 1 |  |  |  |
| *Arhgap32* | GRIT |  |  | 2 |  |
| *Baiap2* | IRSp53 |  | 3 | 4 |  |
| *Cadm1* | SynCam |  | 5 |  |  |
| *Camk2a* | αCaMKII | 6 |  |  |  |
| *Cnksr2* | MAGUIN-1 | 7–9 |  |  |  |
| *Cyfip1* | CYFIP1 | 10,11 | 12 | 13,14 |  |
| *Dlg2* | PSD-93 |  | 15 | 16–19 |  |
| *Dlg3* | SAP102 | 20–23 |  |  | 300189 |
| *Dlg4* | PSD-95-GK | 24 | 25 |  |  |
| *Dlgap1* | GKAP |  |  | 26 |  |
| *Dlgap2* | GKAP2 |  | 27 | 28 |  |
| *Git1* | Git1 | 29 |  | 30 |  |
| *Glul* | Glul | 31 |  |  | 138290 |
| *Gnb1* | Gnb1 | 32 |  |  | 139380 |
| *Gria1* | GluR1 | 33 |  | 34,35 |  |
| *Grin2a* | NR2A-dC | 33,36 |  |  | 138253 |
| *Grin2b* | NR2B-dC | 33,36 |  |  | 138252 |
| *Hras* | H-Ras | 37–40 |  |  | 190020 |
| *Kras* | K-Ras | 41–45 |  |  | 190070 |
| *Nf1* | NF1 | 46–48 |  |  | 162200 |
| *Nsf* | NSF |  |  | 13 |  |
| *Shank2* | SHANK2 | 49 | 27 |  | 603290 |
| *Syngap1* | SynGAP | 33,50–57 | 27,58,59 | 60 | 603384 |
| *Tnik* | TNiK | 61 |  | 62–64 | 610005 |
| *Tspyl2* | CINAP | 65,66 |  |  |  |
